# *In silico* Analysis of Nonsynonymous Genomic Variants within *CCM2* Gene Reaffirm the Existence of Dual Cores within Typical PTB Domain

**DOI:** 10.1101/2021.11.09.467991

**Authors:** Akhil Padarti, Ofek Belkin, Johnathan Abou-Fadel, Jun Zhang

## Abstract

**Purpose:** The objective of this study is to validate the existence of dual cores within the typical phosphotyrosine binding (PTB) domain and to identify potentially damaging and pathogenic nonsynonymous coding single nuclear polymorphisms (nsSNPs) in the canonical PTB domain of the *CCM2* gene that causes cerebral cavernous malformations (CCMs).

**Methods:** The nsSNPs within the coding sequence for PTB domain of human *CCM2* gene, retrieved from exclusive database search, were analyzed for their functional and structural impact using a series of bioinformatic tools. The effects of the mutations on tertiary structure of the PTB domain in human CCM2 protein were predicted to examine the effect of the nsSNPs on tertiary structure on PTB Cores.

**Results:** Our mutation analysis, through alignment of protein structures between wildtype CCM2 and mutant, indicated that the structural impacts of pathogenic nsSNPs is biophysically limited to only the spatially adjacent substituted amino acid site with minimal structural influence on the adjacent core of the PTB domain, suggesting both cores are independently functional and essential for proper CCM2 function.

**Conclusion:** Utilizing a combination of protein conservation and structure-based analysis, we analyzed the structural effects of inherited pathogenic mutations within the CCM2 PTB domain. Our results indicated that the pathogenic amino acid substitutions lead to only subtle changes locally confined to the surrounding tertiary structure of the PTB core within which it resides, while no structural disturbance to the neighboring PTB core was observed, reaffirming the presence of dual functional cores in the PTB domain.

## Introduction

Scaffold proteins have essential roles in various pivotal cellular signaling cascades [1]. One recurring domain shared by scaffolding proteins is the phosphotyrosine binding (PTB) domain [2,3]. As the second-largest family of phosphotyrosine recognition domains, the PTB domain was shown to evolve from pleckstrin homology (PH) domains, as both PTB and PH domains are structurally and functionally comparable sharing the PH superfolds [4]. Contrary to the singular binding pocket in PH domains, the full-length typical PTB domain contains two equal, unique, versatile, and independent binding pockets, or PTB cores, allowing the domain to bind to multiple NPXY motifs present in the cytoplasmic tails of membrane receptors [5]. In this report, we utilized confirmed genetic data to validate the existence of PTB dual cores with biological phenotypes and corresponding genetic data.

CCM2 is a PTB domain-containing protein that binds to CCM1 through either one [6] or two [7] NPXY motifs. CCM1 and CCM3 bind to CCM2 to form the CCM signaling complex that serves as a docking site for other proteins and plays a key role in multiple cellular processes [8]. Several mutations that disrupt the CCM1/CCM2 interaction have been implicated in cerebral cavernous malformations (CCMs). While, it has been experimentally shown that both PTB cores (Core1 and Core2) in CCM2 are independently capable of binding to NPXY motifs on CCM1 [9], the biological relevance on the dual PTB cores remains elusive. To date, more than 150 distinct pathogenic *CCM1/CCM2/CCM3* germline mutations causing CCMs (OMIM 116860) have been implicated in 87-98% of familial CCMs [10]. Approximately 80 nonsynonymous genomic variants have been reported in the *CCM2* gene, and half of them are nonsense mutations or frameshifting-insertions/deletions (INDELs) leading to premature termination codons (PTCs) [11].

Amino acid mutations can affect both the structure and the function of a protein, including post-translational modifications, folding/stability, and ligand binding [12]. Experimental methods for protein structure determination have been established with X-ray crystallography and NMR spectroscopy, generating substantial structural data. With availability of large databases of protein structural information in protein data bank (PDB), various computational approaches have been developed to model the 3D structure of a protein with a known amino acid sequence, known as *in-silico* analysis [13]. Since it is unclear whether both CCM2 PTB cores are necessary and essential for CCM2 functionality, we believe that the effect of a nonsynonymous single nucleotide polymorphism (nsSNP) at the protein level should be studied with *in-silico* methodology to predict the effect of point mutations on PTB domain containing proteins and validate the existence of dual functional PTB cores in CCM2 PTB domain, through phenotype/genotype correlation from defective PTB cores.

## Materials and Methods

### Identification of nonsynonymous genomic variants within the CCM2 gene

To thoroughly investigate the clinical relevance of the nonsynonymous genomic variants within the CCM2 gene, we searched the well-known databases from HGMD (the Human Gene Mutation Database, http://www.hgmd.cf.ac.uk), ClinVar (a public archive with interpretations of clinically relevant variants, http://www.ncbi.nlm.nih.gov/clinvar/), ExAC (the Exome Aggregation Consortium, http://exac.broadinstitute.org), the 1000 Genomes Project (http://www.ncbi.nlm.nih.gov/variation/tools/1000genomes), dbSNP (the Single Nucleotide Polymorphism database, http://www.ncbi.nlm.nih.gov/snp/), OMIM (the Online Mendelian Inheritance in Man, http://www.omim.org), and the Angioma Alliance (https://www.angioma.org/).

### Bioinformatic tools for in-silico analysis

In order to precisely predict the impact of nonsynonymous genomic variants within the CCM2 protein, we utilized well-known bioinformatic tools, which can be categorized into three major groups: protein conservation-based, protein structure-based analysis [14], or a combination thereof, with a selected array of *in-silico* predictive algorithms to evaluate the genomic variants [14,15]. For evolutionary conservative approaches, there are multiple programs such as SIFT (Sorting Intolerant From Tolerant, https://sift.bii.a-star.edu.sg/www/Extended_SIFT_chr_coords_submit.html) [16], PANTHER (Protein ANalysis THrough Evolutionary Relationship, https://www.pantherdb.org/tools) [17], MUTATION ASSESSOR (http://mutationassessor.org/r3/). For homolog modelling (sequence similarities/alignment), we utilized the program PROVEAN (Protein Variation Effect Analyzer, https://provean.jcvi.org/index.php) [18]. For protein structure/function evolutionary conservation-based approach, we selected POLYPHEN-2 (Polymorphism Phenotyping Ver. 2.0, https://genetics.bwh.harvard.edu/pph2/) [19], and MUTATION TESTER (http://www.mutationtaster.org/) [20]. For protein structure stability measurements, we utilized MUPRO (http://mupro.proteomics.ics.uci.edu/), I-MUTANT (http://gpcr2.biocomp.unibo.it/cgi/predictors/I-Mutant3.0/I-Mutant3.0.cgi) [21], HOPE (Have (y)Our Protein Explained, https://www3.cmbi.umcn.nl/hope/method/), and CUPSAT (Cologne University Protein Stability Analysis Tool, http://cupsat.tu-bs.de/) [22] with different parameters to define nonsynonymous variants. The minor allele frequency (MAF) was acquired for each nsSNP from the SNP database. MAF represents the incidence of the gene variant in the general population. We hypothesize if MAF of one nsSNP is less than the overall prevalence of symptomatic CCMs in the general population (0.04%), this nsSNP is likely to be pathogenic.

### Bioinformatic tools for tertiary structural modeling

Currently, there is only one X-ray crystallography structure of PTB domain of CCM2 (binding with an NPXY motif ligand) deposited in PDB (4WJ7) [6]. To better serve our purpose, we utilized MODELLER (https://salilab.org/modeller/9.16/release.html), which uses homology modeling with *ab initio* methods producing solutions that satisfy a set of spatial rules derived from probability density functions and statistical analysis of PTB domain containing protein structures, deposited in PDB [23]. Since, MODELLAR utilizes similar protein structure for structure modelling, the predicted structure was truncated in the C-terminus, since the deposited structure in the PDB of the x-ray crystallographic CCM2 PTB domain is bound to a ligand in the C-terminus. Comparatively, we also used an integrated platform I-TASSER (The iterative threading assembly refinement, https://zhanglab.dcmb.med.umich.edu/I-TASSER/), which is an automated protein structure and functional prediction software based on the sequence-to-structure-to-function paradigm from multiple threading alignments to perform iterative structural assembly simulations [24]. Three-dimensional (3D) atomic models generated by either I-TASSER or MODELLER were then visualized and compared by molecular visualization software, PYMOL (http://www.pymol.org/), CHIMERA (http://www.cgl.ucsf.edu/chimera) [23], and RASWIN (http://www.openrasmol.org/). This process was performed for WT CCM2 and identified CCM2 mutants. The tertiary structure of the two proteins were superimposed and analyzed for any structural differences.

## Results

### Define genomic variants within the PTB domain of CCM2

#### Total number of genomic variants within the PTB domain of CCM2

By searching all available databases, 66 nsSNPs in 49 amino acid positions were identified in the PTB domain of CCM2, in addition two in-frame deletions in exon 2, making a total of 68 nonsynonymous genomic variants within the PTB domain (Supple Table 1). The results of *in-silico* analysis for all nsSNPs are shown (Table 1, Supple Table 1). Among known genetic mutants and our selected candidate mutants, there is significant consistency among predicted results in various *in-silico* analysis; furthermore, the impact of known pathogenic CCM2 missense mutations is reflected by the disrupted functionality of the CCM2 protein (Table 1).

**Table 1:** Pathogenic nsSNPs in both cores of CCM2 PTB domain. The known pathogenic and several potential pathogenic nsSNPs of CCM2 PTB domain are presented. The nsSNPs were chosen based on genetic results from familial CCM cases and high probability of the predicted pathological nature of the mutation. All 66 recorded mutations within CCM2 PTB domain are shown in the supplementary table 1. The secondary structural motif and the core location of each substitution is shown. The SNP nomenclature and MAF for known mutations are also shown if available. Each nsSNP were further evaluated with various *in-silico* tools for pathogenicity. The reference for each reported pathogenic nsSNP in human genetic study evidenced as phenotype/genotype correlation is also provided.

| Mutation | Exon number | PTB Core | Secondary structure | SNP nomenclature | MAF | SIFT |  | MUTATION ASSESSOR |  | PANTHER | CUPSAT |  |  |
| --- | --- | --- | --- | --- | --- | --- | --- | --- | --- | --- | --- | --- | --- |
|  |  |  |  |  |  | Func. Impact | SCORE | Func. Impact | FI score | Func. Impact | Overall Stability | Torsion | ΔΔG (kcal/mol) |
| I76T | 3 | Core1 | β1-α2 | rs756431644 | 8.00E-06 | FUNCTIONAL | 0.03/3.59 | low | 1.1 | probably benign | Destabilising | Unfavourable | -2.54 |
| A98T | 4 | Core1 | α2 | rs780867674 | 1.70E-05 | TOLERATED | 0.06/3.59 | low | 1.79 | probably damaging | Stabilising | Favourable | 2.06 |
| A111P | 4 | Core1 | β2 | rs750889112 | 4.00E-06 | TOLERATED | 0.06/3.59 | medium | 2.2 | probably damaging | Destabilising | Unfavourable | -7.96 |
| L113P | 4 | Core1 | β2 | rs11552377 | 0.06 | FUNCTIONAL | 0.02/3.59 | low | 1.625 | probably benign | Stabilising | Unfavourable | 1.1 |
| L115R | 4 | Core1 | β2 | N/A | N/A | FUNCTIONAL | 0.00/3.59 | medium | 2.72 | probably damaging | Destabilising | Unfavourable | -2.59 |
| V120I | 4 | Core1 | β3 | rs11552377 | 0.175 | TOLERATED | 0.35/3.59 | low | 1.3 | probably benign | Stabilising | Favourable | 0.99 |
| V120D | 4 | Core1 | β3 | rs745788686 | 4.00E-06 | FUNCTIONAL | 0.01/3.59 | medium | 2.2 | probably benign | Destabilising | Favourable | -0.02 |
| A141T | 4 | Core2 | β5 | rs1562908094 | 8.00E-06 | FUNCTIONAL | 0.00/3.59 | medium | 2.54 | probably damaging | Destabilising | Unfavourable | -2.35 |
| R146W | 4 | Core2 | β5 | rs769929401 | 4.80E-05 | FUNCTIONAL | 0.01/3.59 | medium | 2.455 | probably damaging | Destabilising | Unfavourable | -6.53 |
| L152M | 4 | Core2 | β6 | rs760117074 | 8.00E-06 | FUNCTIONAL | 0.00/3.59 | medium | 2.62 | probably damaging | Destabilising | Unfavourable | -2.53 |
| V154G | 4 | Core2 | β6 | rs141353947 | 4.00E-06 | FUNCTIONAL | 0.00/3.59 | medium | 2.6 | probably damaging | Destabilising | Unfavourable | -3.64 |
| L155P | 4 | Core2 | β6 | rs373239614 | 4.00E-06 | FUNCTIONAL | 0.00/3.59 | medium | 2.54 | probably damaging | Stabilising | Unfavourable | 0.87 |
| A179S | 5 | Core2 | α3 | rs373136857 | 1.60E-05 | TOLERATED | 0.39/4.32 | neutral | -0.145 | probably benign | Destabilising | Unfavourable | -3.4 |

| Mutation | MUPRO |  | HOPE | I-MUTANT-2.0 | PROVEAN |  | POLYPHEN-2 | MUTATION TESTER | References |
| --- | --- | --- | --- | --- | --- | --- | --- | --- | --- |
|  | Score | Stability |  |  | Func. Impact | SCORE |  | Func. Impact |  |
| I76T | -2.2330 | Decrease | mutation can disturb this domain and abolish its function | Destabilising | Deleterious | -2.916 | PROBABLY DAMAGING (0.958/0.78/0.95) | Disease-Causing |  |
| A98T | -1.5536 | Decrease | mutation can disturb this domain and abolish its function | Destabilising | Neutral | -0.753 | BENIGN (0.041/0.94/0.83) | POLYMORPHISM |  |
| A111P | -1.3445 | Decrease | might disturb the core structure of this domain. | Stabilising | Deleterious | -2.651 | PROBABLY DAMAGING (0.999/0.14/0.99) | Deleterious | Han et al (2021) |
| L113P | -1.5158 | Decrease | might disturb the core structure of this domain. | Destabilising | Deleterious | -5.043 | likely DAMAGING (0.696/0.86/0.92) | Deleterious | Riant (2013) |
| L115R | -1.2768 | Decrease | residue is located near a highly conserved position | Destabilising | Deleterious | -4.636 | PROBABLY DAMAGING (1.0/0.0/1.0) | Deleterious | Fisher (2015) |
| V120I | -0.7517 | Decrease | POLYMORPHISM | Destabilising | Neutral | 0.262 | BENIGN (0.015/0.96/0.79) | Benign | Pileggi (2010) |
| V120D | -1.7446 | Decrease | might be damaging to the protein and abolish its function | Destabilising | Deleterious | -4.717 | PROBABLY DAMAGING (0.978/0.76/0.96) | Deleterious | Ishii et al (2020) |
| A141T | -1.3542 | Decrease | might be damaging to the protein and abolish its function | Destabilising | Deleterious | -3.034 | Likely DAMAGING (0.628/0.8/0.84) | Deleterious | - |
| R146W | -1.3085 | Decrease | residue is located near a highly conserved position | Destabilising | Deleterious | -5.008 | BENIGN (0.030/0.95/0.82) | Disease-Causing | - |
| L152M | -0.6805 | Decrease | might be damaging to the protein and abolish its function | Destabilising | Neutral | -1.692 | Likely DAMAGIN (0.518/0.82/0.81) | Deleterious | - |
| V154G | -2.4206 | Decrease | might be damaging to the protein and abolish its function | Destabilising | Deleterious | -4.93 | PROBABLY DAMAGING (0.996/0.55/0.98) | Deleterious | - |
| L155P | -2.0019 | Decrease | might be damaging to the protein and abolish its function | Destabilising | Deleterious | -5.376 | PROBABLY DAMAGING (0.996/0.55/0.98) | Deleterious | Fisher (2015) |
| A179S | -0.4483 | Decrease | might be damaging to the protein and abolish its function | Destabilising | Neutral | 0.173 | BENIGN (0.000/1.00/0.00) | POLYMORPHISM |  |

### Genomic variants with in-frame deletions leading to conformational changes in both PTB Cores

#### Large Exon 2 deletion [p.(Pro11_Lys68del)]

This 58 amino acid in-frame deletion was one of the first discovered genomic variants in CCM2 mutation screening [25,26], and has been consistently observed [27,28]. Experimental data showed this in-frame deletion abolishes the interaction between CCM1/CCM2, indicating this portion of the peptide sequence plays an important role in the CCM2/CCM1 interaction [26]. Our analysis indicates that the deletion encompasses a portion of the β1 strand within Core1 (Fig. 1A, Supple Video 1A). β-Sheet 1 contains only 3 β-strands in the deletion mutant, one fewer than the WT. The majority of the PTB domain remains overlapping between the mutant and the WT PTB domain, however, the α3 helix in Core2 is surprisingly distorted indicating that this large deletion results in overwhelming structural changes to both Core1 and Core2 of the CCM2 PTB domain.

**Figure. 1:**
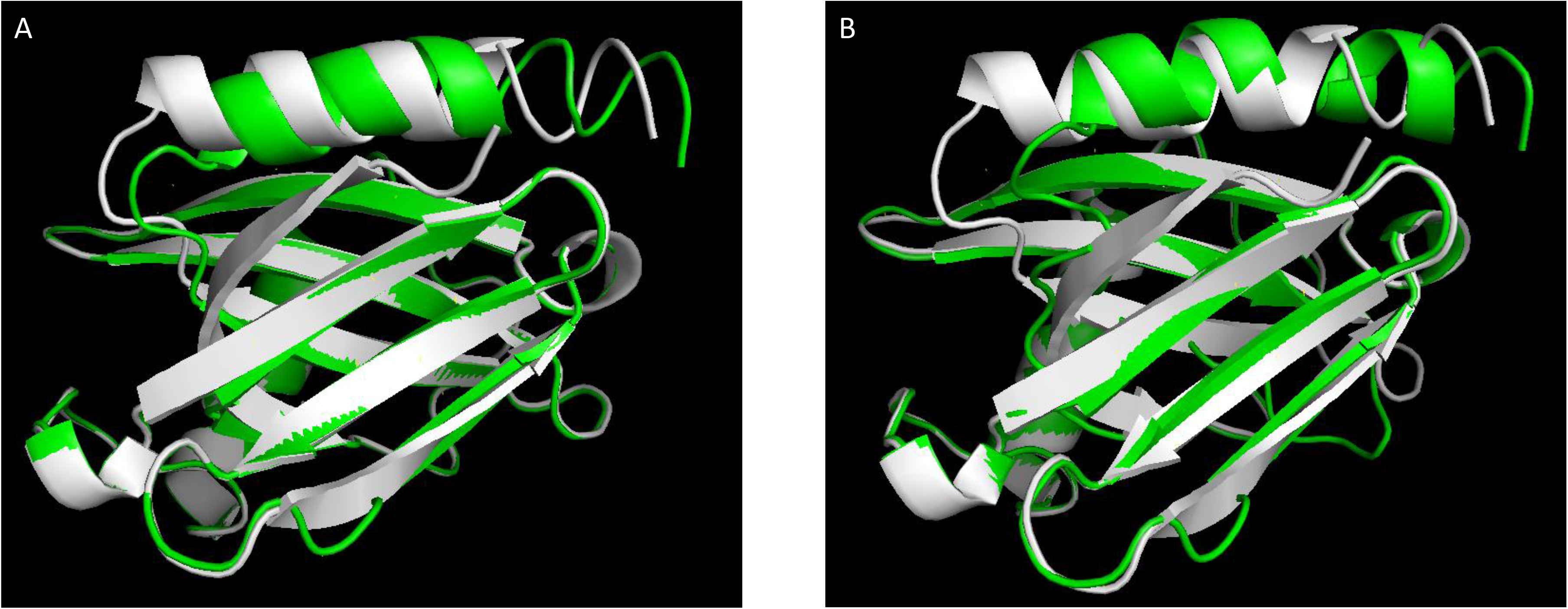
N-terminus in-frame deletion in Core 1 leads to conformational changes in both cores of CCM2 PTB domain. Deletions were modelled by I-TASSER and alignments were visualized using PYMOL. WT CCM2 (white) and mutant (green) are shown. The α3 helix is shown superior and α2 helix is shown inferior, while β-sheet 1 is shown anterior to β-sheet 2. **A)** 58 amino acid deletion in core1 mutant is illustrated. **B)** In-frame 4 amino acid deletion mutant (65-KEVK-68) is illustrated. Although the β-sheet overlaps between the wildtype and mutant, the α3 helix is shown to be mismatched indicating conformational change in the C-terminus.

#### Four in-frame amino acid deletions within Exon 2 [p.(Lys65_Lys68del)]

Another smaller four amino acid in-frame deletion in the same region of the PTB domain was also reported in two Japanese familial CCM cases [29], indicating the importance of this region for the PTB domain. In this deletion, only the N-terminal portion of Core1 is affected with intact upstream flanking sequences. Our analysis indicate that these four amino acids are in the β1 strand between Core1 and Core2 (Fig. 1B, Supple Video 1B). Similarly in the deletion mutant, β-Sheet 1 contains only 3 β-strands, one fewer than the WT, while β-Sheet 2 contains 2 β-strands, two fewer than the WT. While the majority of the PTB domain between the WT and mutant remains aligned, the α3 helix is misaligned in the mutant, similar to the large exon 2 deletion. It is interesting that this four amino acid deletion alters the structure of the β-barrel more severely than the large 58 amino acid deletion, suggesting that these four amino acids may be essential for Core1 function. The structural changes resulting from this deletion encompasses both Core1 and Core 2 of the PTB domain.

### Genomic variants with nsSNPs only have local effects

#### nsSNPs in Core2

CCM disease associated with nsSNPs in Core2, such as L198R and L213P, were among the first discovered genomic variants identified in CCM mutation screening [30–32]. Several nsSNPs (all MAFs < 4e-4, symptomatic CCM incidence in general population) found in Core2 of CCM2 were modelled with I-TASSER and MODELLER (A141T, R146W, L152M, V154G, L155P, A179S) (Fig. 2, Supple Fig 1, Supple Video 2). A141T (Fig. 2, Supple Fig. 1A, Supple Video 2A, 2E) is predicted to be a destabilizing, unfavorable, and rare mutation (MAF < 10e-5). This mutation is in β5 strand, with tertiary structure changes encompassing β5 and a portion of the β5/β6 loop. This is due to the polar side chain of threonine relative to alanine in the WT. L152M (Fig. 2, Supple Fig. 1B, Supple Video 2B, 2F) and V154G (Fig. 2, Supple Fig. 1C, Supple Video 2C, 2G) are two other destabilizing, unfavorable, and rare mutations (MAF< 10e-4) encompassing the β6 strand and these structural perturbations in both mutations are limited to the β6 strand only. The amino acid side chains of both CCM2 mutants are more polar than the side chains of the WT CCM2. L155P (Fig. 2, Supple Fig. 1D, Supple Video 2D, 2H) is a known CCM2 mutation resulting in CCM phenotype [6]. This mutation is in the β6 strand, resulting in structural changes in β6 strand and β6/β7 loop. While leucine and proline both have hydrophobic side chains, proline results in a peptide backbone twist resulting in local structural disturbance. All six Core2 mutations analyzed (A141T, R146W, L152M, V154G, L155P, A179S) show only local structural alterations, but none of these mutations lead to backbone distortion or have structural effects on Core1 of the PTB domain (Fig. 2).

**Figure. 2:**
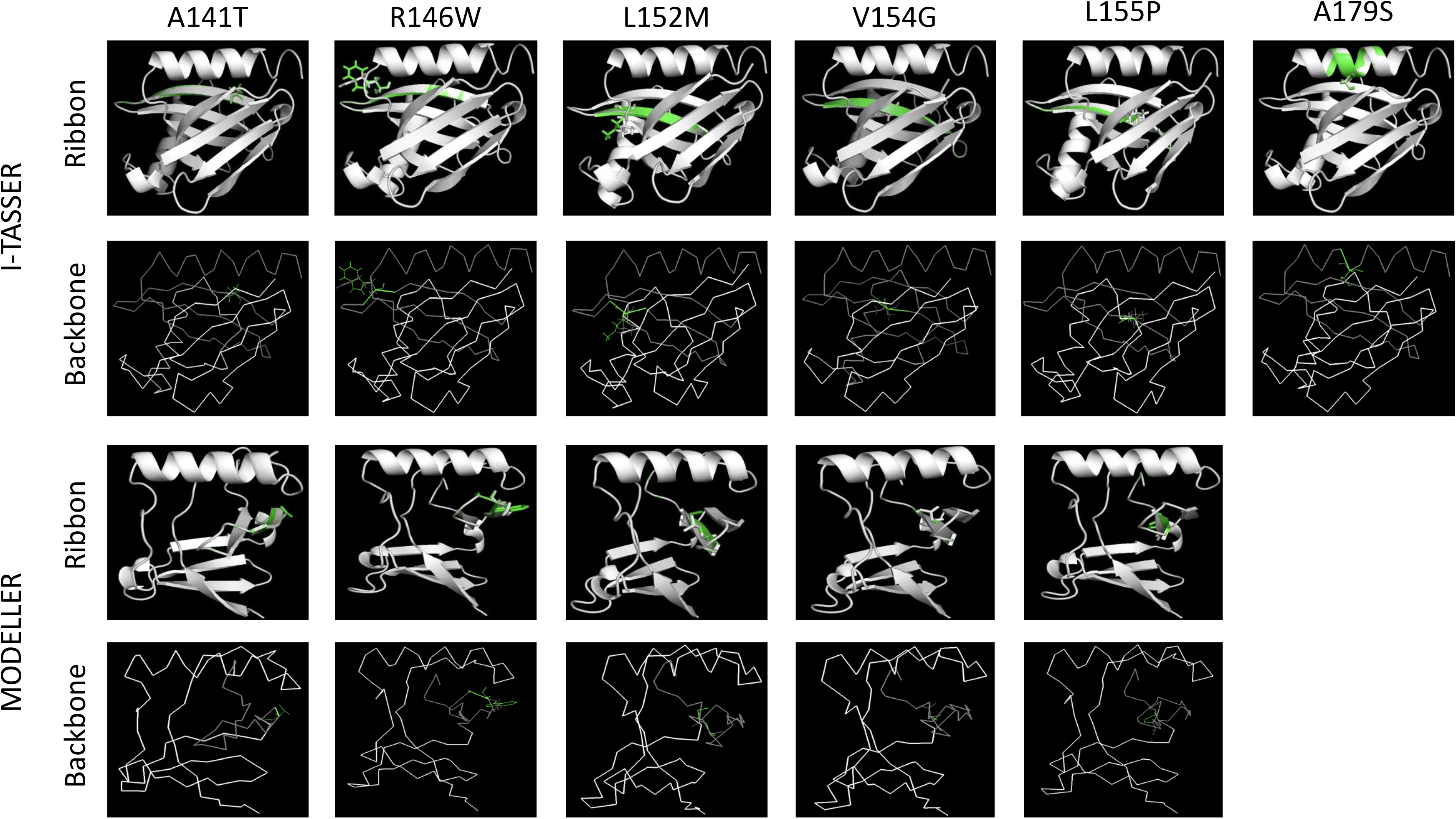
nsSNPs in Core 2 lead to local disturbance of substituted amino acids without perturbing 3D conformation in Core 1 of CCM2 PTB domain in I-TASSER and MODELLER. For I-TASSER 3D structure orientation (Row 1, 2), the α3 helix is shown superior and the α2 helix is shown inferior, while β-sheet 1 is shown anterior to β-sheet 2. For MODELLER 3D structure orientation (Row 3, 4), the α2 helix is shown superior, while the full β-sheet 1 is shown inferior and the partial β-sheet 2 is shown on the right. For the ribbon models (Row 1, 3), the nsSNP mutant (green) is superimposed on the wildtype (white) in the ribbon conformation while the substituted amino acids are shown in stick configuration to demonstrate local conformational change. For the backbone models (Row 2, 4), the same nsSNP is superimposed on the wildtype in backbone conformation with mutant amino acids in line conformation to explore any possible peptide backbone distortion. For the backbone models (Row 2, 4), each core is shown in the different color: wildtype core1 (white), wildtype core2 (gray), mutant core1 (red), and mutant core2 (green) to highlight the two PTB cores (both red and green backbone from mutant can only be visualized if there is a distortion between mutant and wildtype). Several 3D PDB images are also provided as supplements. The nsSNPs are A141T (first column), R146W (second column), L152M (third column), V154G (fourth column), L155P (fifth column), A179S (last column). The C-terminal portion (α3 helix and partial β-sheet 2) of the CCM2 PTB domain model is absent due to the CCM2 x-ray crystallographic structure data in the PDB database which emphasizes ligand binding in the C-terminal PTB core2 and for the same reason, A179S mutant is unable to be generated with MODELLER. Only subtle local disturbance was seen surrounding the substituted amino acids and no amino acid backbone distortion was observed.

#### nsSNPs in Core1

More pathogenic nsSNPs have been reported in the N-terminus of the PTB domain corresponding to Core1 than Core2. Several nsSNPs in Core1 have been modelled with I-TASSER and MODELLER (I76T, A98T, A111P, L113P, L115R, V120D) (Fig. 3, Supple Fig. 2, Supple Video 3). A111P (Fig. 3, Supple Fig. 2A, Supple Video 3A, 3E) is a pathogenic mutation in the β2 strand (β-sheet 1), with minimal overlap between the amino acid side chains in both tertiary structure modalities [33]. There is misalignment in the β2, β2/3 loop, and α1/β2 loop between the two structures. Proline is more sterically bulky than alanine and possesses a larger local twist in the peptide backbone, resulting in a shortened β2 strand. L113P (Fig. 3, Supple Fig. 2B, Supple Video 3B, 3F) is a known mutation that results in familial CCMs [32] that is present in β2 strand, with structural perturbations in β1/β2 strands and β2/ β3 loop. The side chain of proline results in a backbone peptide chain twist resulting in delayed formation of β2 strand. L115R (Fig. 3, Supple Fig. 2C, Supple Video 3C, 3G), a known pathological CCM mutation [6], resides in the β2 strand, with all structural differences limited to β2 strand and the β2/3 loop. Arginine has a charged side chain and all hydrophobic interactions with WT leucine are severed. V120D (Fig. 3, Supple Fig. 2D, Supple Video 3D, 3H), a known mutation resulting in a Japanese familial CCM pedigree [34], is located in the β3 strand, with all structural differences limited to the β3 strand and β3/4 loop. Similar to L115R, V120D results in inhibition of all hydrophobic interactions of valine with the charged amino acid side chain of aspartate. In sum, similar to Core2 nsSNPs mutations, all six mutations within Core1 (I76T, A98T, A111P, L113P, L115R, V120D) show only local structural alterations, but none of these mutations lead to backbone distortion or have structural disruption on Core2 of the PTB domain (Fig. 3).

**Figure. 3:**
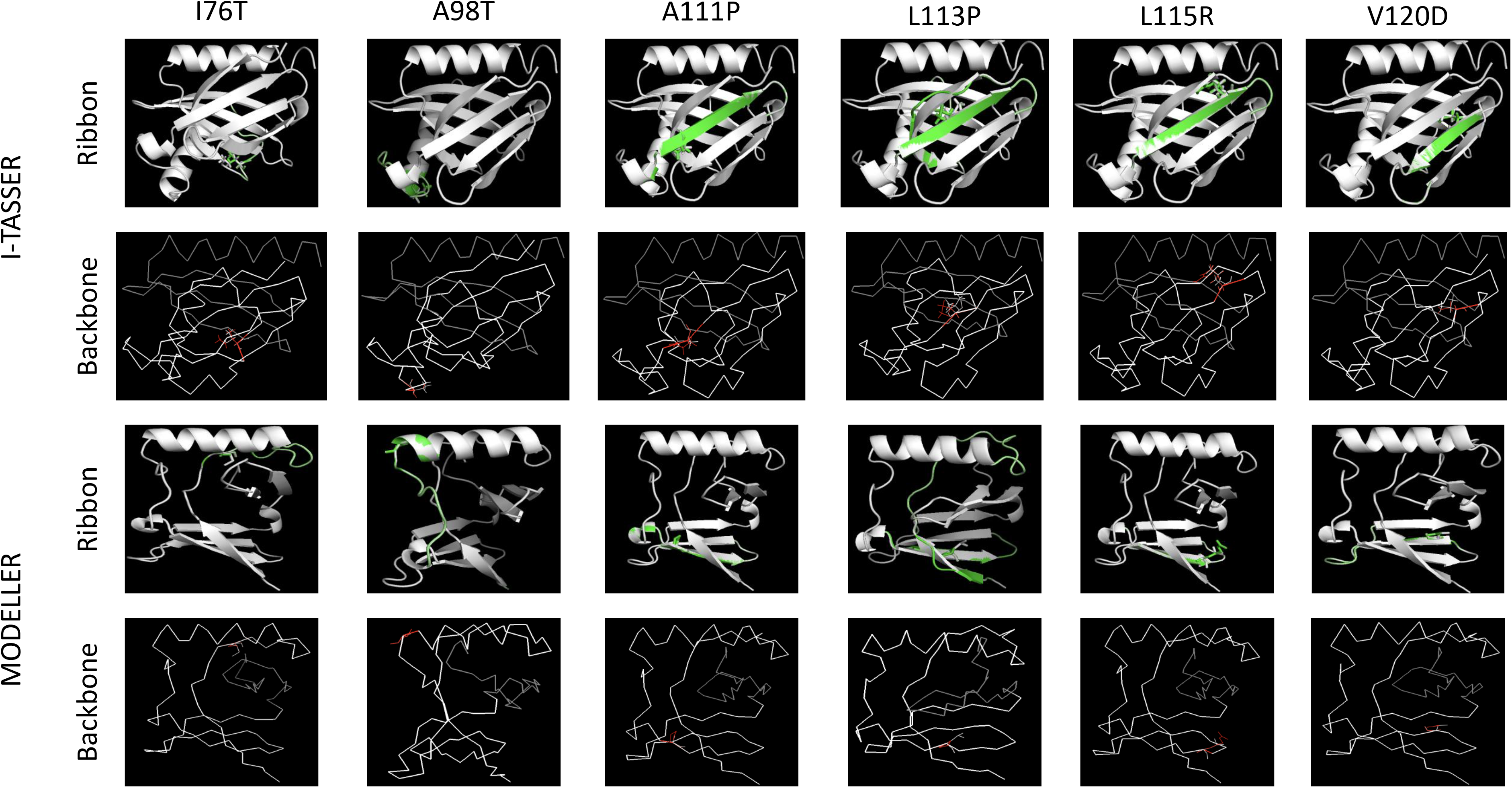
nsSNPs in Core 1 lead to local disturbance of substituted amino acids without perturbing 3D conformation in Core 2 of CCM2 PTB domain in I-TASSER and MODELLER. Diagram layout is similar to Figure 2, I-TASSER (Row 1, 2) and MODELLER (Row 3, 4) 3D structure orientation, ribbon (Row 1, 3) and backbone (Row 2, 4) are displayed in the same fashion. The nsSNPs are I76T (first column), A98T (second column), A111P (third column), L113P (fourth column), L115R (fifth column), V120D (last column). Similar to Core2 mutants, only subtle local disturbance was seen surrounding the substituted amino acids and no amino acid backbone distortion was observed.

#### In-silico analysis of a common genomic variant within the CCM2 PTB domain questions its role as causative mutation

It is interesting to note that a common nsSNP (rs11552377) has been one of the most widely reported *CCM2* mutations in *CCM2* gene screening analysis as the genetic cause of CCMs [27,34–41]. This nsSNP was predicted to be in the “benign” end of the CCM disease phenotype spectrum [36], with data suggesting this nsSNP affects splicing [38]. However, this variant is quite common (MAF range: 0.13-0.17) (Table 1). Furthermore, the predicted possible alterations of amino acid substitution (V120I) generated were consistently defined as benign/polymorphic (Table 1), challenging previous mutation reports to be spurious. Our *in-silico* analysis indicates V120I mutation is in the β3 strand in β-sheet 1 of Core1 (Supple Fig. 3, 4, Supple Video 4). The associated tertiary structure perturbation is limited to β3 strand and β3/4 loop, which is the more flexible region of the PTB domain. The side chains of both valine and isoleucine are overlapping, similar sterically and equally hydrophobic, maintaining the hydrophobic side chain interactions. Therefore, we conclude this mutation (V120I) is polymorphic and not pathogenic. Further mutation screening in newly described novel coding exons of *CCM2* [9] might be warranted for these familial CCM cases.

## Discussion

### In-silico analysis of genetic mutations among CCM genes

Since identification of causative genes of CCMs, there have been many attempts to utilize *in-silico* analysis with bioinformatics tools to interpret the genetic variants identified within CCM genes [27,28,32,38,39,42–45]. However, the majority of these targeted genetic variants among three known *CCM* genes are either nonsense mutations or frame-shift mutations, making the outcomes of the *in-silico* analysis irrelevant to protein structural investigations. To date, one protein tertiary structure of CCM2 PTB domain binding to NPXY motif has been deposited in PDB (4WJ7), determined by x-ray crystallography [6]. Although this work was primarily focused on the traditionally recognized PTB functional pocket, Core2, it provided the foundation for one of our protein structural simulation programs, probability density functions-based MODELLER [23], making our novel integrated *in-silico*/*structural simulation* analysis more thorough. Furthermore, recent efforts for *in-silico* analysis of Core1 in a genetic variant that is co-segregated in a large Chinese CCM pedigree [33] provide us with additional strong evidence for supporting this methodology. On the structural level, each amino acid substitution frequently resulted in perturbations within the tertiary structure of the PTB domain. These perturbations were due to disruption of the electrostatic/hydrophobic interactions, yet the structural alteration was limited to the surrounding region of the substitution within the PTB domain. The two in-frame deletions are the exception to this observance. Both in-frame N-terminal deletions resulted in alteration of Core1 and Core2, strongly suggesting that the β1 strand/α2 helix at the N terminus may be equally essential for maintaining structure stability of the entire PTB domain as the C terminus, in contrast with previous reports [6,46]. Our data conclude that nsSNPs in Core1 did not disrupt Core2 and vice versa, indicating that both binding pockets have independent functional roles in their interactions with NPXY motifs, and dysfunction of either binding pocket is sufficient to initiate pathogenesis of CCMs.

#### In-silico analysis provides novel candidates for future CCM2 mutational screening and assists in excluding nsSNPs as causal mutations

A significant portion of the CCM patients have no known causative mutations identified [38]. It may be difficult to differentiate between non-pathogenic and pathogenic nsSNPs in these patients. *In-silico* methodology can be used to identify high-risk nsSNPs as potential causal mutations for these patients while ruling out certain nsSNPs as benign polymorphisms. Our *in-silico* analysis revealed that one missense mutation, V120I, currently considered a causal mutation of CCMs, is in fact a relatively common polymorphism and unlikely to result in phenotype. Similarly, this methodology also helped us to identify several potential candidates pathological nsSNPs along with known pathogenic nsSNPs reported previously (Table 1). Those candidate pathogenic nsSNPs can be further evaluated through MAF and their performance in *in-silico* analytical tools. In sum, our integrated *in-silico* analysis will be useful in future CCM mutation screening to identify pathogenic mutations and excluding normal variants (Suppl. Table 1).

## Conclusion

This analysis demonstrated that both PTB-cores in CCM2 have independent functional binding pockets and mutations in either one can result in CCM phenotype without disrupting the conformation of the neighboring core, validating our PTB dual core theory. One important limitation to our methodology is analyzing the extent of cooperative binding between the two PTB cores. It is rather likely that cooperative binding between the two PTB cores exists optimizing the binding ability of the CCM2 PTB domain. However, despite *in silico* analysis is a proven valid and robust methodology for analyzing protein domains, the extent of this cooperative binding cannot be evaluated. Future efforts are needed to further explore the dual core nature of the PTB domains and their potential unique cellular partners for each binding pocket.

## Abbreviations

PTB: phosphotyrosine binding
PH: pleckstrin homology
CSC: CCM signaling complex
CCMs: cerebral cavernous malformations
PDB: protein data bank
nsSNP: nonsynonymous single nucleotide polymorphism
WT: wildtype
INDELs: insertions/deletions
PTCs: premature termination codons
SIFT: Sorting Intolerant From Tolerant
PANTHER: Protein ANalysis THrough Evolutionary Relationship
PROVEAN: Protein Variation Effect Analyzer
POLYPHEN-2: Polymorphism Phenotyping
HOPE: Have (y)Our Protein Explained
CUPSAT: Cologne University Protein Stability Analysis Tool
MAF: minor allele frequency
I-TASSER: the iterative threading assembly refinement

## Supplementary Introduction

In the early practice of *in-silico* analysis, inconsistent results were usually encountered from different modeling and algorithms, preventing any conclusions to be made about the true impact of these nsSNPs. However, the current approaches of combining several *in silico* methodologies makes *in-silico* analysis a powerful and robust tool to evaluate the functional and structural effects of amino acid mutations. Structural basis for pathogenic mutations of CCM2 has been explored only in the context of protein structural biology, or in the nature of genetic mutations. Furthermore, there is lack of an *in-silico* analysis on the functional and structural impacts of pathogenic nsSNPs of the CCM2 PTB domain at the protein level. In this report, we utilized methods combining both physical force fields and free parameters fitted with experimental data to predict the structural impacts of novel missense variants identified in familial CCM patients. This is the first study combining missense mutations in human genetic studies with *in-silico* analysis and comparative structural evaluations to spatially analyze the impact of potentially damaging pathogenic nsSNPs in the *CCM2* gene. The objective of this study is to utilize the most reliable bioinformatics tools to assess any tertiary structural changes with known pathogenic CCM2 variants to investigate the requirement of whether one or both functional cores within the CCM2 PTB domain are needed.

### Supplement Legend

**Supplementary Figure 1:**
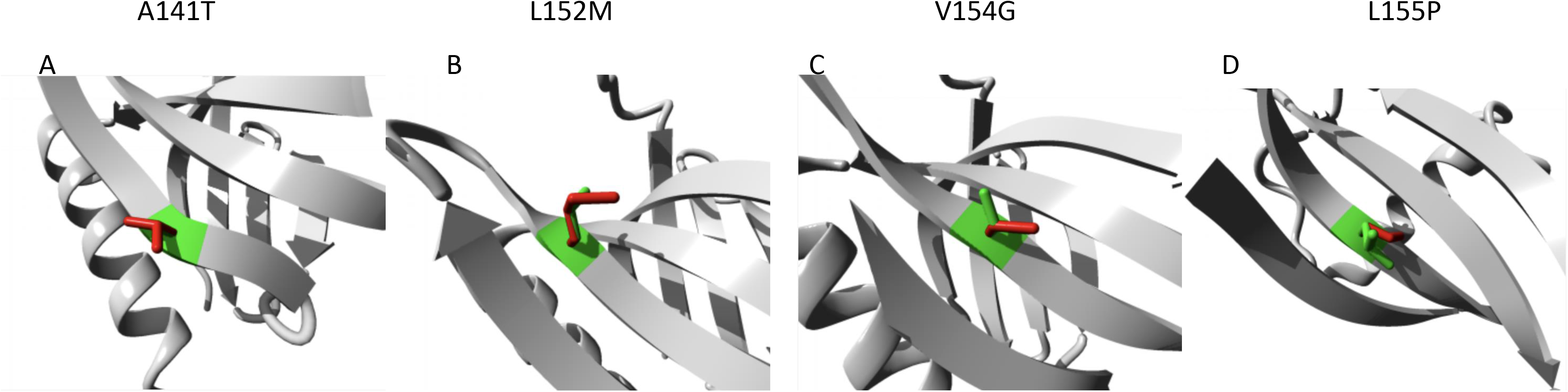
Local disturbances caused by a single amino acid substitution coded by nsSNPs in Core2 of CCM2 PTB domain. The local amino acid differences between the WT and mutant in Core2 mutations are detailed (HOPE). The WT (green) and mutant (red) amino acids are shown in stick configuration. **A)** A141T mutant superimposed by WT CCM2. Alanine is a hydrophobic residue relative to the polar amino acid threonine. **B)** L152M mutant superimposed by WT CCM2. Leucine is a hydrophobic residue relative to the polar amino acid side chain of methionine. **C)** V154G mutant superimposed by WT CCM2. Valine is a sterically bulky residue relative to glycine lacking a peptide side chain. **D)** L155P mutant superimposed by WT CCM2. Leucine and proline are both sterically bulky hydrophobic chains, but proline is an aromatic amino acid that results in twisting of the peptide backbone.

**Supplementary Figure 2:**
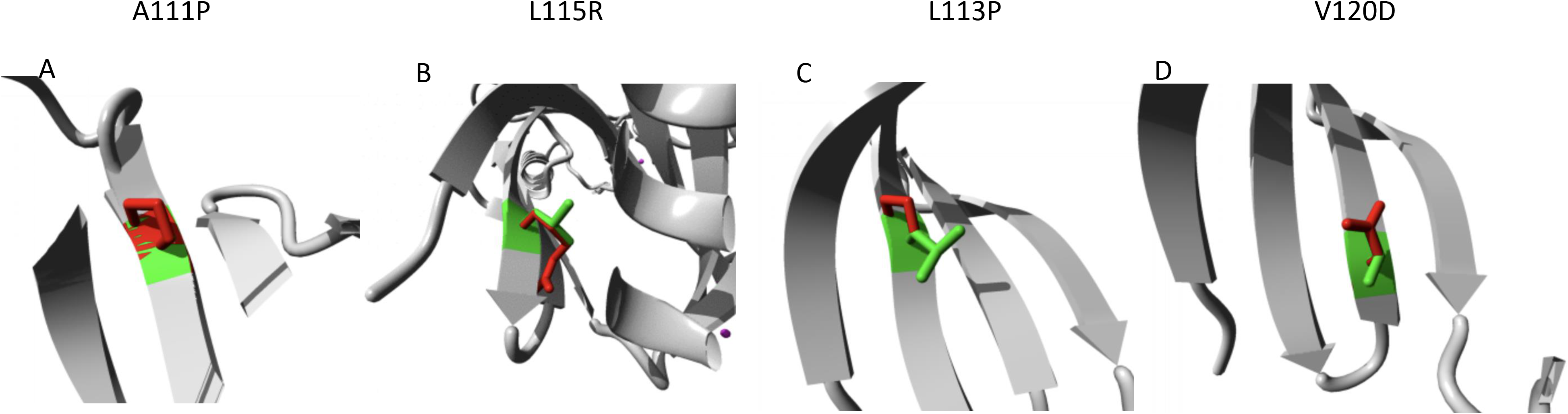
Local disturbances caused by a single amino acid substitution coded by nsSNPs in Core1 of CCM2 PTB domain. The local amino acid differences between the WT and mutant in Core2 mutations are detailed (HOPE). The WT (green) and mutant (red) amino acids are shown in stick configuration. **A)** A111P mutant superimposed by WT CCM2. Alanine is a small hydrophobic amino acid residue while proline is a sterically bulky amino acid residue. **B)** L113P mutant superimposed by WT CCM2. Both leucine and proline have hydrophobic side chains, but proline is a ringed amino acid that results in a peptide backbone twist. **C)** L115R mutant superimposed by WT CCM2. Leucine is a hydrophobic residue while arginine is a charged amino acid. **D)** V120D mutant superimposed by WT CCM2. Valine is a hydrophobic amino acid while aspartate is charged.

**Supplementary Figure 3:**
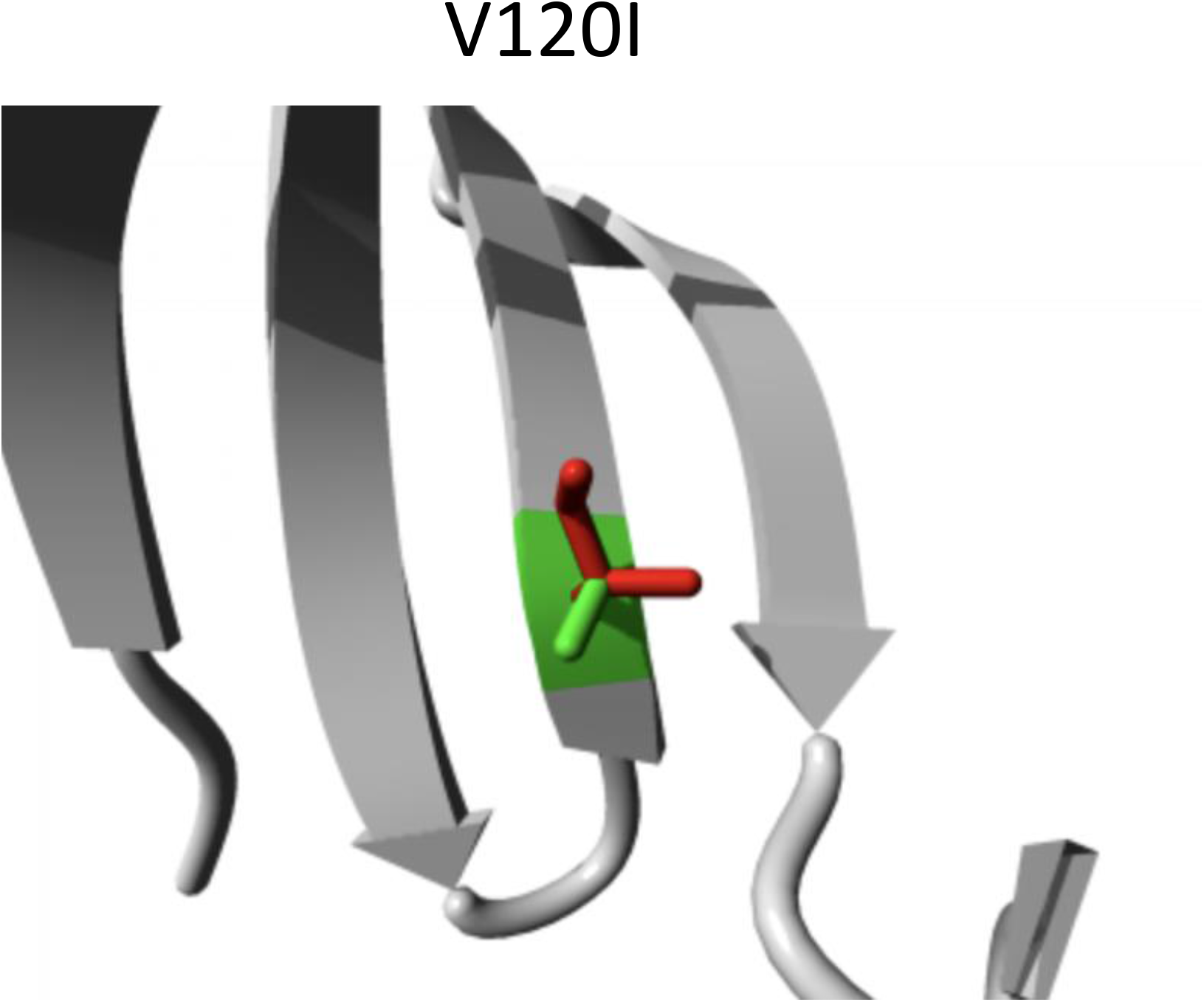
No major local disturbances were observed with V120I substitution. The local amino acid differences between WT and mutant Core1 mutations are detailed (HOPE). The WT (green) and mutant (red) amino acids are shown in stick configuration. V120I mutant superimposed by WT CCM2. Both Isoleucine and valine and bulky amino acid side chains with similar steric and electronic effects.

**Supplementary Figure 4:**
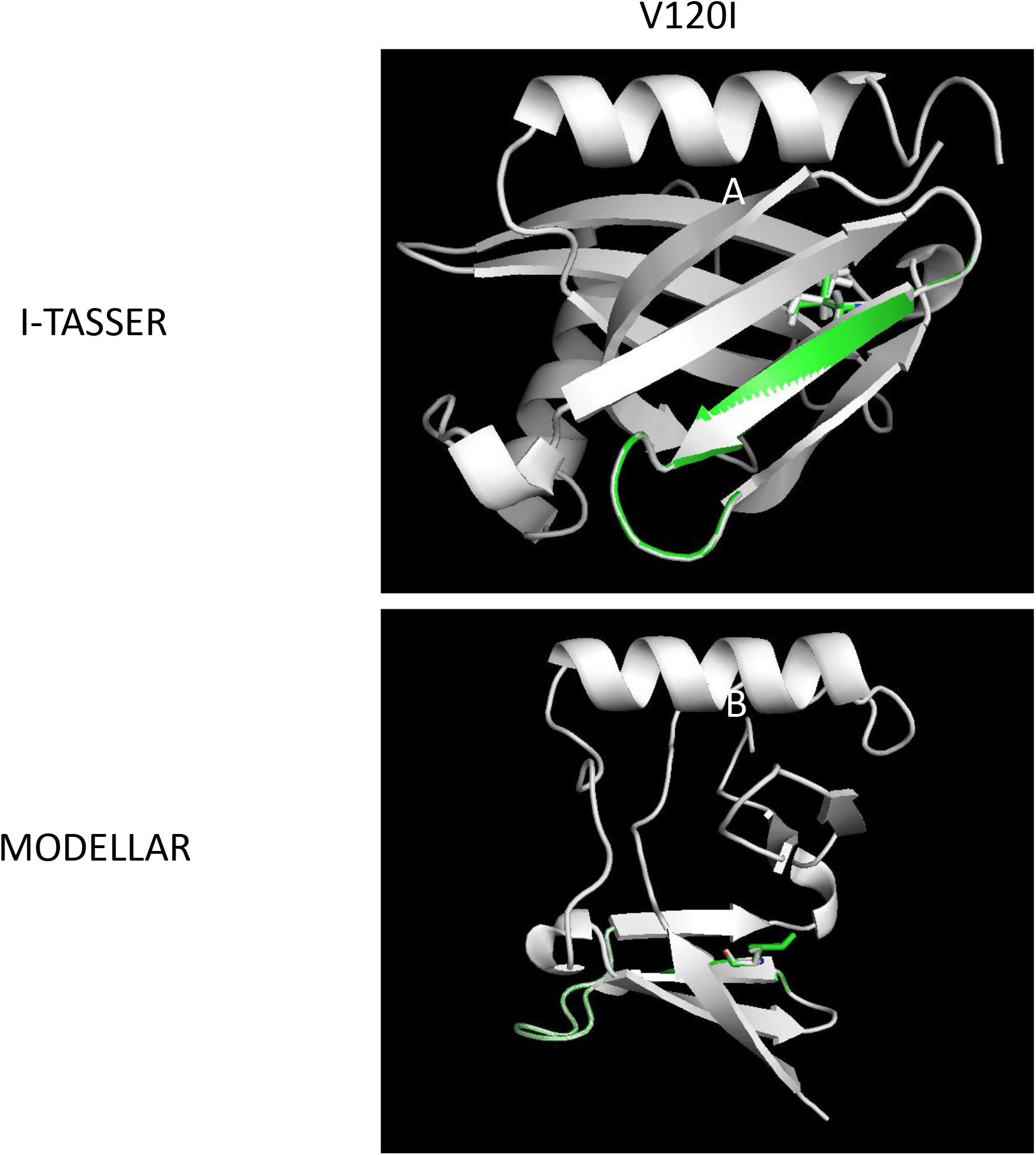
The V120I substitution matches well with WT 3D structure. V120I substitution was modelled by I-TASSER/MODELLER and superimposition between WT (white) and mutants (green) were visualized with PYMOL. For 3D structure orientation, in the ITASSER predictions, the α3 helix is shown superior and the α2 helix is shown inferior, while β-sheet 1 is shown anterior to β-sheet 2; in the MODELLER predictions, the α2 helix is shown superior, while the full β-sheet 1 is shown inferior and the partial β-sheet 2 is shown on the right. Mutant amino acids are shown in stick configuration. The 3D PDB images are also provided as supplement. A) V120I mutant superimposed by WT CCM2 via I-TASSER. B) V120I mutant superimposed by WT CCM2 via MODELLER.

### Supplementary Videos

**Supplementary Video 1: 3D Rotating in-frame deletion in Core 1 mutants of CCM2 PTB domain.** Deletions were modelled by I-TASSER/MODELLAR and alignment was visualized with PYMOL. The 3D rotating structures are shown as videos with superimposition of WT CCM2 (white) and mutant (green) figures. Mutant amino acids are shown in stick configuration. **A)** 28 amino acid deletion in core1 mutant shown. **B)** In-frame 68-KEVK-65 deletion mutant is shown.

**Supplementary Video 2: 3D Rotating nsSNPs in Core 2 of CCM2 PTB domain.** Amino acid substitutions coded by nsSNPs were modelled by I-TASSER/MODELLAR and alignments were visualized with PYMOL. The 3D rotating structures are shown as videos with superimposition of WT CCM2 (white) and mutant (green) figures. Mutant amino acids are shown in stick configuration. **A)** A141T mutant superimposed by WT CCM2 via I-TASSER. **B)** L152M mutant superimposed by WT CCM2 via I-TASSER. **C)** V154G mutant superimposed by WT CCM2 via I-TASSER. **D)** L155P mutant superimposed by WT CCM2 via I-TASSER. **E)** A141T mutant superimposed by WT CCM2 via MODELLAR. **F)** L152M mutant superimposed by WT CCM2 via MODELLAR. **G)** V154G mutant superimposed by WT CCM2 via MODELLAR. **H)** L155P mutant superimposed by WT CCM2 via MODELLAR.

**Supplementary Video 3: 3D Rotating nsSNPs in Core 1 of CCM2 PTB domain.** Amino acid substitutions coded by nsSNPs were modelled by I-TASSER/MODELLAR and alignment was visualized with PYMOL. The 3D rotating structures are shown as videos with superimposition of WT CCM2 (white) and mutant (green) figures. Mutant amino acids are shown in stick configuration. **A)** A111P mutant superimposed by WT CCM2 via I-TASSER. **B)** L113P mutant superimposed by WT CCM2 via I-TASSER. **C)** L115R mutant superimposed by WT CCM2 via I-TASSER. **D)** V120D mutant superimposed by WT CCM2 via I-TASSER. **E)** A111P mutant superimposed by WT CCM2 via MODELLAR. **F)** L113P mutant superimposed by WT CCM2 via MODELLAR. **G)** L115R mutant superimposed by WT CCM2 via MODELLAR. **H)** V120D mutant superimposed by WT CCM2 via MODELLAR.

**Supplementary Video 4: 3D Rotating V120I Substitution in Core 1 of CCM2 PTB domain.** V120I Substitution was modelled by I-TASSER/MODELLAR and alignment was visualized with PYMOL. The 3D rotating structure is shown as a video with superimposition of WT CCM2 (white) and mutant (green) figures. Mutant amino acids are shown in stick configuration. **A)** V120I mutant superimposed by WT CCM2 via I-TASSER. **B)** V120I mutant superimposed by WT CCM2 via MODELLAR.

### Supplementary Table

**Supplementary Table 1: All nsSNPs identified in CCM2 PTB domain.** All 66 nsSNPs identified through our database search are shown with the amino acid position. The secondary structural motif and the core location of each substitution is shown. If the amino acid is found between two secondary structures, then both structural moieties are outlined with a hyphen. The SNP nomenclature and MAF for known mutations are also shown if available. Each nsSNP were further evaluated with seven representative *in-silico* tools for pathogenicity.

